# Nerve injury converts Schwann cells in a persisting repair state in human neuroma tissue

**DOI:** 10.1101/2024.05.13.593891

**Authors:** Stefanie Deininger, Jakob Schumacher, Anna Blechschmidt, Jialei Song, Claudia Klugmann, Gregor Antoniadis, Maria Pedro, Bernd Knöll, Sofia Meyer zu Reckendorf

## Abstract

Peripheral nerve injury (PNI) induces neuroma formation at the severed nerve stump resulting in impaired nerve regeneration and functional recovery in patients. So far, molecular mechanisms and cell types present in the neuroma impeding on regeneration have only sparsely been analyzed. Herein we compare resected human neuroma tissue with intact donor nerves from the same patient. Neuroma from several post-injury timepoints (1-13 months) were included, thereby allowing for temporal correlation with molecular and cellular processes. We observed reduced axonal area and percentage of myelin producing Schwann cells (SCs) compared to intact nerves. However, total SOX10 positive SC numbers were comparable. Notably, markers for SCs in a repair mode including c-JUN and SHH (sonic hedgehog) and SC proliferation (pH3) were upregulated in neuroma, suggesting presence of SCs in repair rather than differentiated status. In agreement, in neuroma, pro-regenerative markers such as phosphorylated i.e. activated CREB (pCREB), ATF3, GAP43 and SCG10 were upregulated. Neuroma tissue was infiltrated by several types of macrophages. Finally, when taken in culture, neuroma SCs were indistinguishable from controls SCs with regard to proliferation and morphology. However, cultured neuroma SCs retained a different molecular signature from control SCs including increased inflammation and reduced gene expression for differentiation markers such as myelin genes.

In summary, human neuroma tissue consists of SCs with a repair status and is infiltrated strongly by several types of macrophages.

## Introduction

In the human peripheral nervous system (PNS), severed axons have the potential for limited regeneration. So far, most molecular and cellular mechanisms associated with successful axon regeneration have been deciphered in rodent models. These rodent models identified the denervated Schwann cell (SC) as key driver for nerve regeneration in addition to neuron intrinsic growth promoting mechanisms. For this, myelinating SCs undergo de-or trans-differentiation to switch into a pro-regenerative repair SC phenotype (Bosch-Queralt et al., 2023; Jessen and Mirsky, 2019). In the mouse, the latter is accompanied by upregulation of molecular repair markers including the transcription factor c-JUN (Arthur-Farraj et al., 2012; Gomez-Sanchez et al., 2015). Indeed, c-Jun is functionally involved in PNS regeneration since conditional c-Jun ablation in mice results in impaired repair cell formation (Arthur-Farraj *et al*., 2012). These repair SCs comprise a state of high plasticity as indicated by elevated proliferation rates and elongation to form guiding tracks for regenerating axons called Büngner bands (Jessen and Mirsky, 2021). After successful axon re-growth, repair SCs re-differentiate into myelinating SCs accompanied by upregulation of different molecular markers. This includes EGR2 (Zhang et al., 2023) whose deficiency in the mouse PNS prevents SCs from differentiation to myelinating SCs (Topilko et al., 1994). In general, the regenerative response of human nerves after injury is incomplete without surgical interference compared to rodent models with almost full functional recovery (Barnes et al., 2022; Scheib and Hoke, 2013). So far, molecular and cellular responses accounting for this difference are poorly understood. In injured human nerves, neuroma formation at the nerve stump (“stump neuroma”) or neuroma-in-continuity (NIC) occurs (Huang et al., 2023; Yang et al., 2022) which rarely happens spontaneously in rodent PNI. In humans, such neuroma tissue develops over months after injury and is removed by surgical intervention. Neuroma tissues consist of SCs and disorganized sprouting axons. Furthermore, few available reports show enrichment of fibrotic material and inflammatory cells (Durrenberger et al., 2006; Karsy et al., 2018; Mahan et al., 2019; Vora et al., 2007). Such scar formation in injured nerves is considered to impede nerve regeneration as shown in rodent models (Atkins et al., 2006). On the molecular level, a faster lipid metabolism shut-down most likely resulting in more efficient conversion to repair SCs in rodent compared to human nerves was described (Meyer Zu Reckendorf et al., 2020). In addition, upregulation of molecules associated with repulsive axon guidance such as Semaphorins and WNTs was described in neuroma (Tannemaat et al., 2007; van Vliet et al., 2021). Besides differences, few data available also point at conserved responses between human and rodent PNI since human SCs also adopt a c-JUN positive repair SC identity (Wilcox et al., 2020). For instance, c-JUN was not found in uninjured nerves whereas higher abundance was reported in traumatic neuromas and even more c-JUN positive SCs were found in schwannoma (Shivane et al., 2013).

In this study we provide a comprehensive histological and molecular analysis of human neuroma tissue by comparing responses to non-injured intact sural nerves derived from the same patient. We analyzed nerves in terms of axonal density, regeneration markers, myelination, inflammation and SC phenotype. Finally, we investigated molecular and morphological SC changes between neuroma and control SCs in primary cultures.

Taken together, we identified a repair SC phenotype which suggests a regenerative potential of these SCs to myelinate injured axons even months after trauma. Further we suggest that these repair SCs are localized in a highly inflammatory and fibrotic milieu impeding on axonal growth.

## Materials and Methods

### Patient recruitment and sample preparation

Neuroma tissue and sural nerves were derived from peripheral nerve transplantation surgeries performed at the Department of Neurosurgery, section of peripheral nerve repair at the district hospital of Günzburg (Ulm University, Germany). According to the study protocol, written informed consent was given by all patients before surgery. As “neuroma age” we define the time span between injury and surgical removal. Here, it should be kept in mind that neuroma formation might not exactly start right after injury but could develop later. Included patients were ≥ 18 years. All surgery was performed under general anesthesia. Experiments were approved by the local ethical review board (Nr. 91/16, 261/20 and 224/22).

Resected neuroma tissue was stratified for the different experiments as depicted in Table 1. Functional recovery after PNI depends on neuroma age and is typically poor if neuroma persist for more than one year. In order to analyze changes associated with neuroma age, neuroma tissue was collected at timepoints ranging from one month up to 8 months for the majority of cases post injury (Table 1). As control, we employed sural nerve pieces which were mostly derived from the same patients. These sural nerve pieces were remaining nerve parts not required for nerve transplantation surgeries to bridge the nerve stumps after neuroma removal.

**Table 1:** Patient information.

| # | Age/<br>gender | Injury<br>type | Nerve<br>type | Surgical<br>procedure | Neuroma<br>type | Latency<br>(days) | Sample used for |
| --- | --- | --- | --- | --- | --- | --- | --- |
| 1 | 43/f | iatrogenic | SN | Split repair | NIC | 281 | H (S+N) |
| 2 | 50/m | mc accident | BP | Tx | n.a. | 79 | H (S+N) |
| 3 | 57/f | iatrogenic | RN | Tx | stump | 54 | H (S+N) |
| 4 | 36/m | mc accident | BP | Tx | NIC | 203 | H (S+N) |
| 5 | 62/f | iatrogenic | BP | Tx | NIC | 234 | H (S+N) |
| 6 | 51/m | bike accident | BP | Tx | n.a. | 262 | H (S+N) |
| 7 | 61/m | cut injury | MN | Tx | Stump | 41 | H (S+N) |
| 8 | 41/m | Impact injury | UN | Split repair | NIC | 84 | H (S+N) |
| 9 | 41/m | stab injury | BP | Tx | Stump | 57 | H (N) |
| 10 | 56/m | bike accident | BP | Tx | Stump | 179 | H (N) |
| 11 | 28/m | iatrogenic | PN | Tx | Stump | 119 | H (S+N), iDISCO (N) |
| 12 | 30/m | mc accident | BP | Tx | Stump | 161 | H (S+N), iDISCO (S+N) |
| 13 | 63/m | shoulder disloc. | BP | Tx | NIC | 131 | H (S+N), iDISCO (S+N) |
| 14 | 60/m | mc accident | BP | Tx | NIC | 173 | H (S+N), iDISCO (N) |
| 15 | 25/m | train accident | BP | Tx | NIC | 189 | H (N), iDISCO (S+N) |
| 16 | 46/f | knee dislocation | PN | Tx | NIC | 163 | H (S+N) |
| 17 | 19/m | cut injury | UN | Tx | Stump | 32 | H (S+N) |
| 18 | 24/m | mc accident | BP | Tx | Stump | 118 | H (N) |
| 19 | 72/m | shard injury | SN | Tx | Stump | 27 | H (S+N) |
| 20 | 79/m | shoulder disloc. | BP | Tx | NIC | 178 | H (S+N) |
| 21 | 55/m | iatrogenic | MN | Tx | NIC | 155 | H (S+N) |
| 22 | 46/f | iatrogenic | MN | Tx | NIC | 155 | H (S+N) |
| 23 | 66/f | humerus fract. | MN | Tx | NIC | 392 | iDISCO (N) |
| 24 | 32/m | iatrogenic | PN | Tx | Stump | 174 | H (S+N), qPCR (S+N) |
| 25 | 57/m | chain saw acc. | RN | Tx | Stump | 248 | H (S+N), qPCR (S+N) |
| 26 | 18/m | knee dislocation | PN | Tx | NIC | 197 | qPCR (S+N) |
| 27 | 34/f | knee dislocation | PN | Tx | NIC | 150 | qPCR (S+N), ICC (S+N) |
| 28 | 66/f | iatrogenic | RN | Tx | NIC | 162 | qPCR (S+N), ICC (S+N) |
| 29 | 62/m | mc accident | BP | Tx | Stump | 191 | ICC (S+N) |
| 30 | 56/m | iatrogenic | UN | Split Repair | NIC | 216 | ICC (S+N) |
| 31 | 46/m | iatrogenic | RN | Split repair | NIC | 85 | qPCR (S+N) |
| 32 | 43/m | flex saw injury | MN | Tx | Stump | 155 | qPCR (S+N) |
| 33 | 27/m | cut injury | PN | Tx | Stump | 365 | qPCR (S+N) |
| 34 | 51/m | stab injury | BP | Tx | Stump | 193 | qPCR (S+N) |

Abbreviations: BP, brachial plexus; f, female; H, histology; ICC, immunocytochemistry; m, male; MN, median nerve; mc, motor cycle; N, neuroma; n.a., not assessed; NIC, neuroma in continuity; PN, peroneal nerve; qPCR, qPCR from cultured Schwann cells; RN, radial nerve; S, sural nerve; SN, sciatic nerve; TX, transplantation; UN, ulnar nerve

Samples from 34 patients (34 neuroma and 30 intact sural control nerves) were included in this study. The majority of patients were males (26/34; Table 1). The mean patient age at the time of surgery was 47.1 years (± 15.5 years; min.: 18 years, max.: 79 years). The average time between injury and surgery was 164.8 days (± 83.3 days; min.: 27 days, max.: 392 days). The majority of neuromas was derived from the brachial plexus (14/34) followed by peroneal (6/34) and median nerve (5/34), radial (4/34), ulnar (3/34) and sciatic nerve (2/34).

Samples (neuroma and sural nerve) for RNA isolation were frozen immediately at - 80°C. Samples for histology were fixed in 4% FA (formaldehyde) over-night. Nerves assigned for cell culture were kept in Ringer solution at 37 °C until subsequent treatment (maximum of three hours).

### Histology and Immunocytochemistry

Sural nerves and neuroma were embedded in paraffin after fixation (4% paraformaldehyde, overnight at 4°C) and 5 μm microtome sections were prepared. Primary antibodies used were as follows: anti-βIII tubulin (rabbit, 1:5000, BioLegend, 801201; mouse, 1:3000, BioLegend, 801213), anti-MBP (mouse, 1:2000, BioLegend, 836504), anti-pCREB (rabbit, 1:800, Cell Signaling, 9198), anti-c-JUN (rabbit, 1:500, Cell signaling, #9165), anti-S100ß (rabbit, 1:1000, Abcam, ab52642), anti-S100ß (mouse, 1:500, invitrogen, XA3457432), anti-CD11c (D3V1E) XP® (rabbit, 1:400, Cell signaling, #45581), anti-CD163 (D6U1J; rabbit, 1:1000, Cell signaling, #93498), anti-GAP43 (rabbit, 1:1000, Chemicon, AB5220), anti-SHH (mouse, 1:15, clone 5E1, RRID: AB_2188307), anti-pHistone H3 (rabbit, 1:500, abcam, ab177218), anti-IBA1 (rabbit, 1:1000, WACO, 019-19741), anti-CD68 (D4B9C) XP® (rabbit, 1:500, Cell signalling, #76437), anti-Ki67 (rabbit, 1:400, Thermo Fisher, MA5-14520), anti-SOX10 (mouse, 1:100, abcam, ab218522), anti-SCG10 (“Stathmin-2/STMN2” rabbit, 1:1000 Novus Biologicals, NBP1-49461), anti-BDNF (rabbit, 1:250, santa cruz, sc-546), anti-CD45 (rabbit, 1:1000, abcam, ab10558), anti-SMA (mouse, 1:200, Millipore, CBL171), anti-ATF3 (rabbit, 1:500, Novus biologicals, NBP1-85816).

Immunohistochemistry was performed using biotin-conjugated secondary antibodies (1:500; BA-1000, Vectorlabs) and a peroxidase-based detection system using the ABC kit (PK-6100, Vectorlabs) and DAB as substrate. Alternatively, anti-mouse or rabbit Alexa 488 and Alexa 546 conjugated secondary antibodies (1:500, Thermo Fisher Scientific, A-11003, A-11008/A-11001/A-11003/A-11071/A-11039) were used. Phalloidin staining of SCs in culture was performed by incubation of TexasRed Phalloidin (1:100, Biotium, 00033) for one hour at room temperature.

#### *iDISCO* (immunolabeling-enabled three-dimensional imaging of solvent-cleared organs)

iDISCO^+^ procedures have followed a standard protocol (https://idisco.info/idisco-protocol/). In brief, human neuroma and sural nerve samples were cross-sectionally trimmed into 3mm-thick pieces. Samples were dehydrated in methanol series and bleached in 5% H_2_O_2_. After rehydration, immunolabeling was performed at 37℃ as follows: 3d in permeabilization solution, 3d in blocking solution, 7d in primary antibodies, washed overnight, 7d in secondary antibodies and washed overnight. Samples were dehydrated in methanol series, incubated in 66% dichloromethane/33% methanol for 3h at RT and finally cleared in ethyl cinnamate (Sigma 112372). Primary antibodies used were as follows: anti-SMA (mouse, 1:200, Millipore, CBL171), anti-βIII tubulin (rabbit, 1:2000, BioLegend, 801201). Anti-mouse Alexa 488 or rabbit Alexa 546 conjugated secondary antibodies (1:500, Thermo Fisher Scientific, A-11003, A-11001) were used.

#### Imaging and data processing of iDISCO^+^

Cleared tissue was imaged with Ultra Microscope II light-sheet microscopy from LaVision with a 6.3x objective. Samples were fixed with UV glue in a stage chamber filled with ethyl cinnamate. ImSpector Pro Software (LaVision BioTec) was used to operate the microscopy with appropriate laser strength and sheet width to achieve the best signal-noise ratio. Images were taken at a resolution of 2048x2048 pixels with a 1 μm step between z-stacks. Images were taken in fascicles of sural nerve and nerve-rich area of neuroma recognized by βIII tubulin signal. Images were processed with Imaris 9.9.1(Oxford Instruments). In brief, z-stack images went through built-in Image Proc program using Gaussian Filter and Background Subtraction to reduce the background signal before reconstruction and analysis. Background noise was further decreased with *Surfaces* function by setting signal outside surface as a 0 intensity. Cavities inside smooth muscle actin (SMA) signals were recognized by *Cells* function and filled by masks in Imaris in order to calculate the true volume of blood vessels. “Filaments” function was used to recognize the blood vessel structure of SMA. SMA signal were quantified using built-in Volume statistics tool. All data were normalized to total volume of βIII tubulin signal.

### Imaging quantification

Quantification of histological and cytochemical fluorescent and bright field images was performed using the ImageJ or QuPath software. For each staining, a threshold was used to set a constant brightness or intensity threshold to define specific staining as compared to background.

Depending on the staining either the number or the area (as indicated in each graph) was quantified by the automated “analyze particles” function of ImageJ. For quantification of axons, nuclei or Schwann cells, the ImageJ function “find maxima” was used. For markers of inflammation (Fig. 5) and SCG10 (Fig. 4G-H) the software QuPath was used for analysis. Here, the functions “pixel classification” or “cell detection” were used to quantify percentage of stained area or number of positive particles respectively. At least two sections per sample were analyzed and the mean value was used for quantification.

The SC area of cultured human SCs was calculated by surrounding the cell body with a “segmented line” in ImageJ and automatically calculating the area out of the measured perimeter.

### Electron microscopy (EM)

Nerves and neuromas were fixed overnight in 4% PFA, post-fixed in 2.5% glutaraldehyde for at least 24 h and embedded in epoxy embedding medium. Semi-thin sections (300 nm) were prepared, stained with toluidine by applying a 0.05% toluidine solution for 10-20 sec and ultrathin sections (80 nm) were prepared within the appropriate area. Axonal diameter and g-ratio was calculated using the perimeters of axon and myelin. The g-ratio was calculated using the formula: g-ratio = diameter_axon_/diameter_fiber_. For each patient, 30-40 random axons and myelin sheaths in three frames were quantified. The mean of each patient was calculated for interindividual comparison.

### Quantitative polymerase chain reaction (qPCR)

RNA was isolated from samples using TRIzol (Qiagen) and the RNeasy kit (Qiagen) according to the manufacturers protocol. Reverse transcription was performed with 1 μg RNA using reverse transcriptase (Promega) and random hexamers for cDNA synthesis. qPCR was performed on a Light Cycler 96 system (Roche) with the TB Green Premix Ex Taq PCR master mix (Takara). For each sample, the LightCycler 96 software detects the threshold cycle value (Ct value). Gene Expression was calculated in relation to RNA levels of the house keeping gene *HPRT* (hypoxanthine phosphoribosyltransferase 1) to account for potential variations in total mRNA amounts. Primers used were as follows:

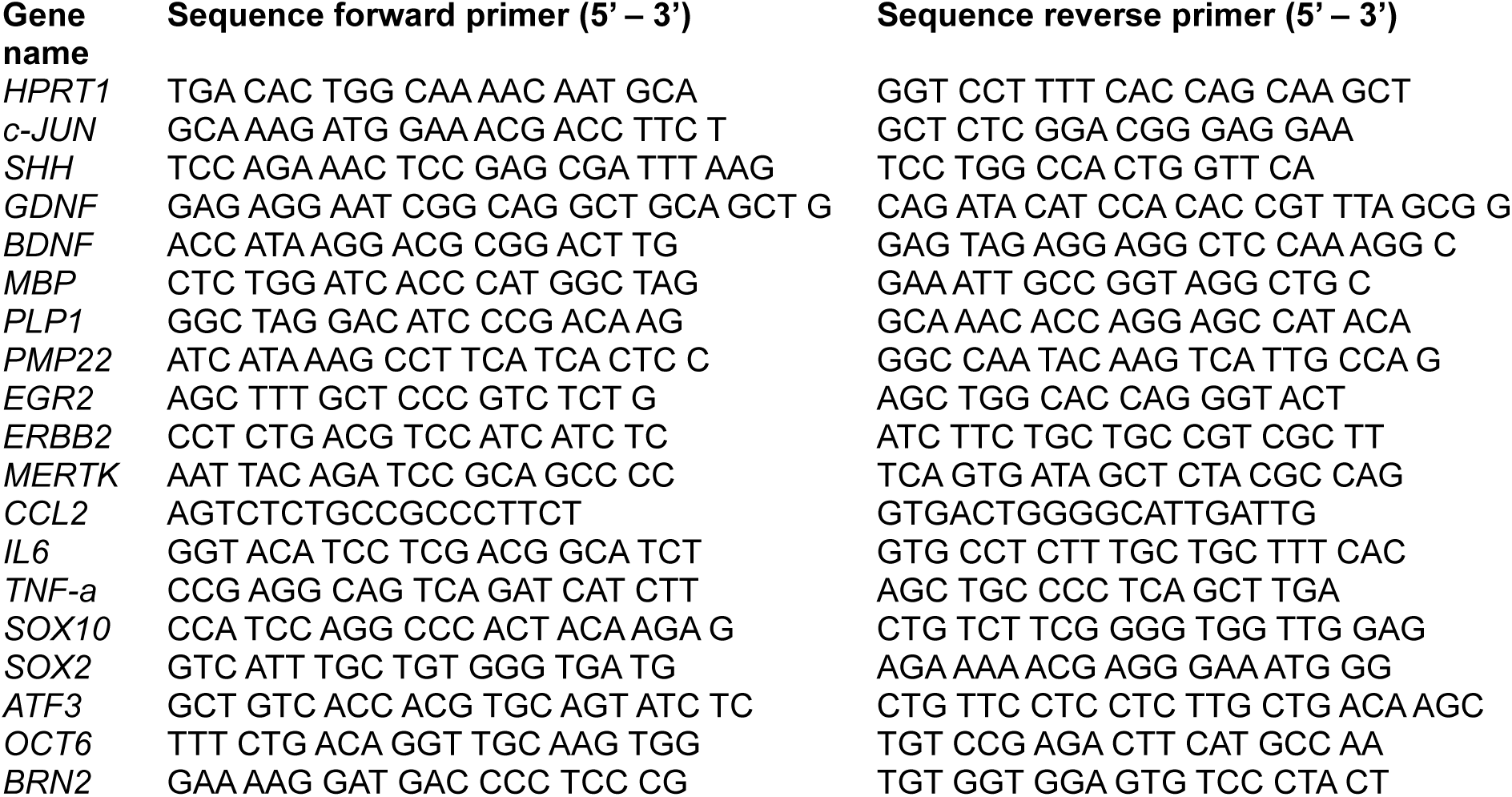

### Primary human SC culture

Epineurium free sural nerves and neuromas were cut in pieces (< 1cm) and transferred into DMEM containing 10% FCS and 1% Pen/Strep for eight to ten days. Medium was changed three times/week. Subsequent enzymatic digestion was done according to the protocol of Haastert et al. (Haastert et al., 2007) followed by trituration. The achieved single cell suspension was sorted in a six well ultra-low attachment plate (Corning Costar®, 3471) over a period of 48-72 hours. SCs were then seeded on poly-L-lysine/laminin coated coverslips or petri dishes and kept in the above-mentioned medium containing recombinant human neuregulin1-ß1/heregulin1-ß1 (0.1 mg/ml, R&D systems, 396HB) for 48 hours. For immunocytochemistry 2x10^3^ cells per coverslip (12 mm Ø) were seeded, for RNA isolation, 4-6x10^5^ cells per well in 6-well plates. Purity of S100β positive cells was ≥ 95% in all evaluated cultures.

### Statistical analysis

At least three biological replicates were analyzed for all experiments. Numbers (n) are indicated in each figure or figure legend. Statistical analysis of data was done with GraphPad Prism software (GraphPad Software, Inc.). Gaussian distribution was tested using the D’Agostino-Pearson omnibus normality test. Since some of the groups were not normally distributed, the non-parametric paired Wilcoxon test (two-sided) was chosen if not mentioned otherwise in the figure legend to compare between sural nerve and neuroma. Further, Pearson’s correlation was used to investigate the correlations between neuroma age and staining intensity for the various parameters. Statistical significance is provided as *, **, *** quoting P ≤ 0.05, 0.01 and 0.001, respectively. SD is provided if not mentioned otherwise.

## Results

### Pronounced axonal loss correlates with neuroma age

In the first set of experiments, the extent of axonal demise in neuroma was analyzed over post-injury time (Fig. 1). Sural control nerves presented a typical axonal organization in fascicles interspersed with connective tissue (Fig. 1A). Neuroma which was surgically removed at four months post injury revealed a similar organization (Fig. 1B). In contrast, in neuroma tissue at later time-points (e.g. up to 8 months) organization in fascicles was diminished (Fig.1C; quantified in D). Nevertheless, even in these long-term neuroma “mini-fascicles” (Fig. 1F) and contained similar axon numbers as sural nerves (Fig. 1E). Quantification of axon number/area irrespective of neuroma age revealed a 20 % reduction in neuroma compared to control nerve (Fig. 1G). Nevertheless, with increasing neuroma age, a decline in βIII tubulin positive axons in the entire nerve was noticed (Fig. 1D) in line with previous reports (Domer et al., 2018; Ebenezer et al., 2007; Wilcox *et al*., 2020).

**Figure 1.**
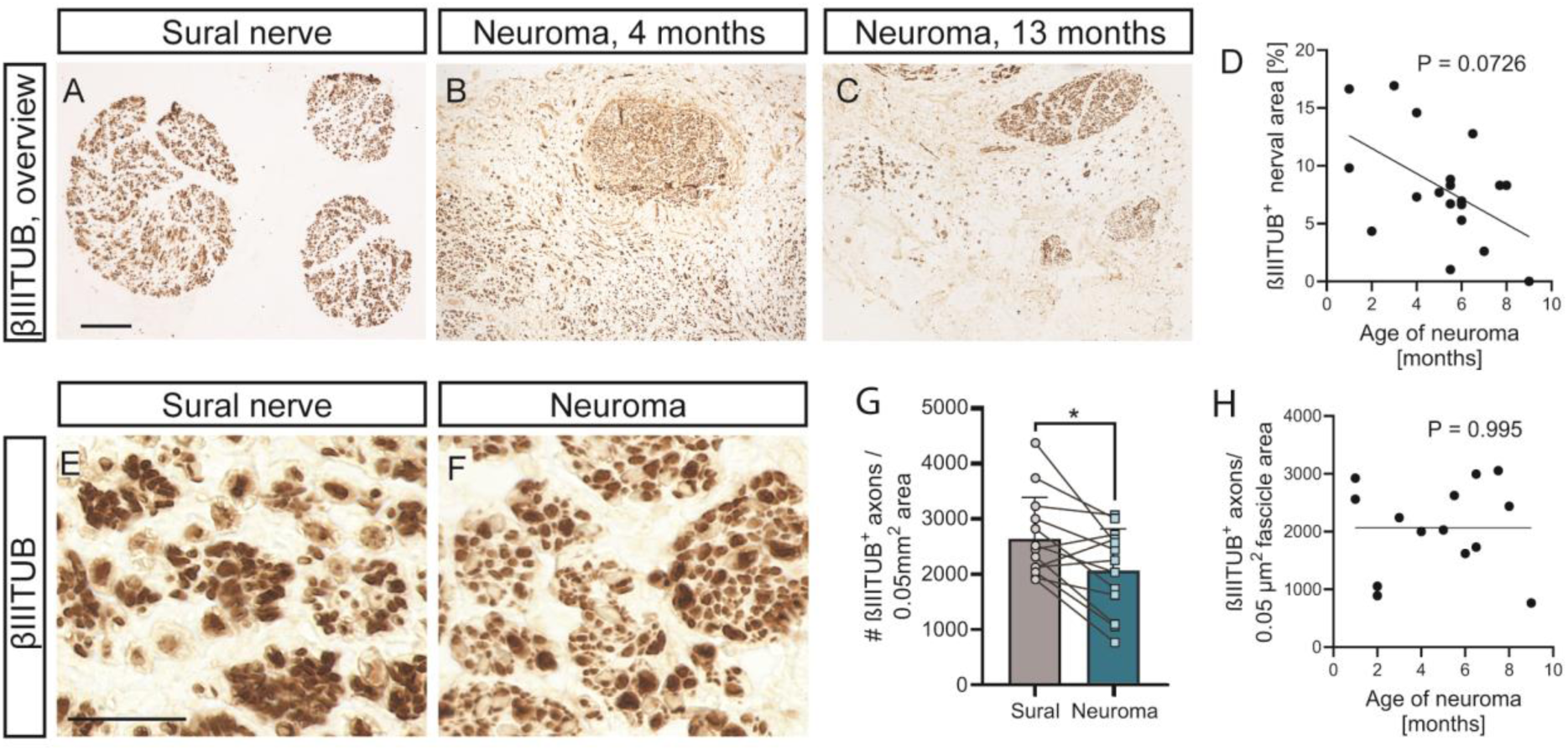
Increased neuroma age correlates with enhanced axonal demise. (A-D) Sural nerve (A) and neuroma tissue (B, C) were stained for axons with βIII tubulin. In freshly developed neuroma (B, four months after injury) axonal organization in fascicles is almost similar to sural control nerves from the same patient. Compared to control nerve (A) and younger neuroma (B), older neuroma clearly showed axonal demise (C) which significantly correlated with age (D). (E-H) If fascicles were still present in neuroma (F), axon density within the fascicle was similar to sural nerve (E). Combining all neuroma ages, axon number within fascicles was decreased by 20% in neuroma compared to control nerve (G) but did not change with neuroma age (H). Each line connects neuroma and sural nerve from the same patient (G). Each black circle in (D, H) depicts one patient. Data are presented as mean ± SD. For statistical analysis, Wilcoxon test was performed in (G), and Pearson’s test in (D, H). Statistical significance is provided as *, **, *** reflecting P ≤ 0.05, 0.01 and 0.001, respectively. Scale bar (A-C) = 100 μm, (E, F) = 25 μm

Besides axonal organization, course and abundance of blood vessels and formation of fibrotic tissue were investigated (Fig. 2). Alpha smooth muscle actin (SMA) is mainly expressed in vascular smooth muscle cells but also can be expressed in fibrogenic cells such as myofibroblasts which contribute to fibrosis. We noticed that neuroma tissue had enhanced presence of SMA positive cells (Fig. 2B, D) in relation to sural nerves where almost no SMA positive cells were detectable in the connective tissue around the fascicles (Fig. 2A, C; quantified in E). This indicates that fibrosis was enhanced in neuroma and quantification revealed persisting fibrosis over neuroma age (Fig. 2F). SMA also labelled nerve resident blood vessels. Here, we used the iDISCO tissue clearing technique to provide 3D spatial reconstruction of axonal and blood vessel trajectories (Fig. 2G-K). In sural nerves, only thin blood vessels were found within the endoneurium of nerve fascicles and most blood vessels were localized in the perineurium (Fig. 2G, I, Supp. Video 1). Also, in sural nerves many blood vessels were mainly running longitudinally alongside the axons (Fig. 2G, I; Supp. Video 1). In contrast, in neuroma, where fascicle structures were less defined (see Fig. 1C), blood vessels and axons were more disorganized. Here, axons were typically intermingled with blood vessels and were frequently crossing each other (Fig. 2H, J; Supp. Video 2). In addition, the abundance of blood vessels was increased in neuroma compared to sural nerve (Fig. 2K).

**Figure 2.**
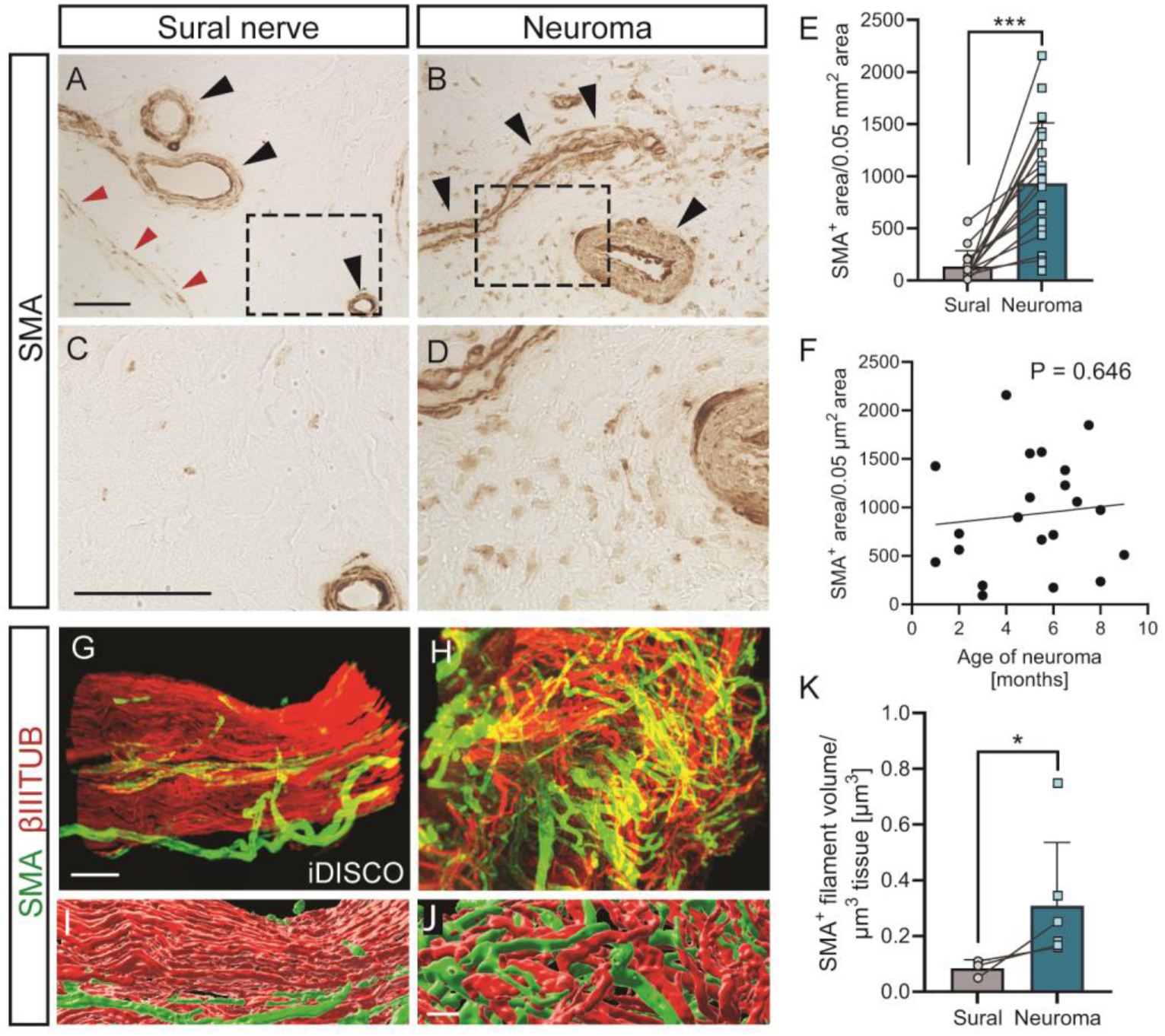
Increased SMA abundance in neuroma tissue. (A-D) SMA positive cells were upregulated in neuroma (B, D) compared to sural nerve (A, C; quantified in E). SMA positive cells were rather constant over neuroma age (F). C, D represent higher magnifications of A and B, respectively. Black arrowheads point at blood vessels, red arrowheads denote a fascicle border (G-K) iDISCO of nerve samples. SMA positive blood vessels (green) and axons (red) were labelled in sural nerve (G, I) and neuroma (H, J). In neuroma, axons and blood vessels were intermingled and disorganized (H, J). Quantification revealed higher abundance of blood vessels in neuroma (K). Each black circle in (F) depicts one patient. Data are presented as mean ± SD. For statistical analysis, Wilcoxon test was performed in (E), and Pearson’s test in (F). Statistical significance is provided as *, **, *** reflecting P ≤ 0.05, 0.01 and 0.001, respectively. Scale bar (A, B) = 25 μm, (C, D) = 25 μm, (G, H) = 50 μm, (I, J) = 20 μm

Taken together, neuroma shows progressive axon loss, enhanced fibrosis and disturbed organization of blood vessels.

### Myelinated axons decrease with increased neuroma age

Above, reduced axonal density was noticed in aged neuroma (Fig. 1). Since axons and SCs form a functional unit, consequences on axon myelination and numbers of myelin producing SCs were assessed (Fig. 3).

**Figure 3.**
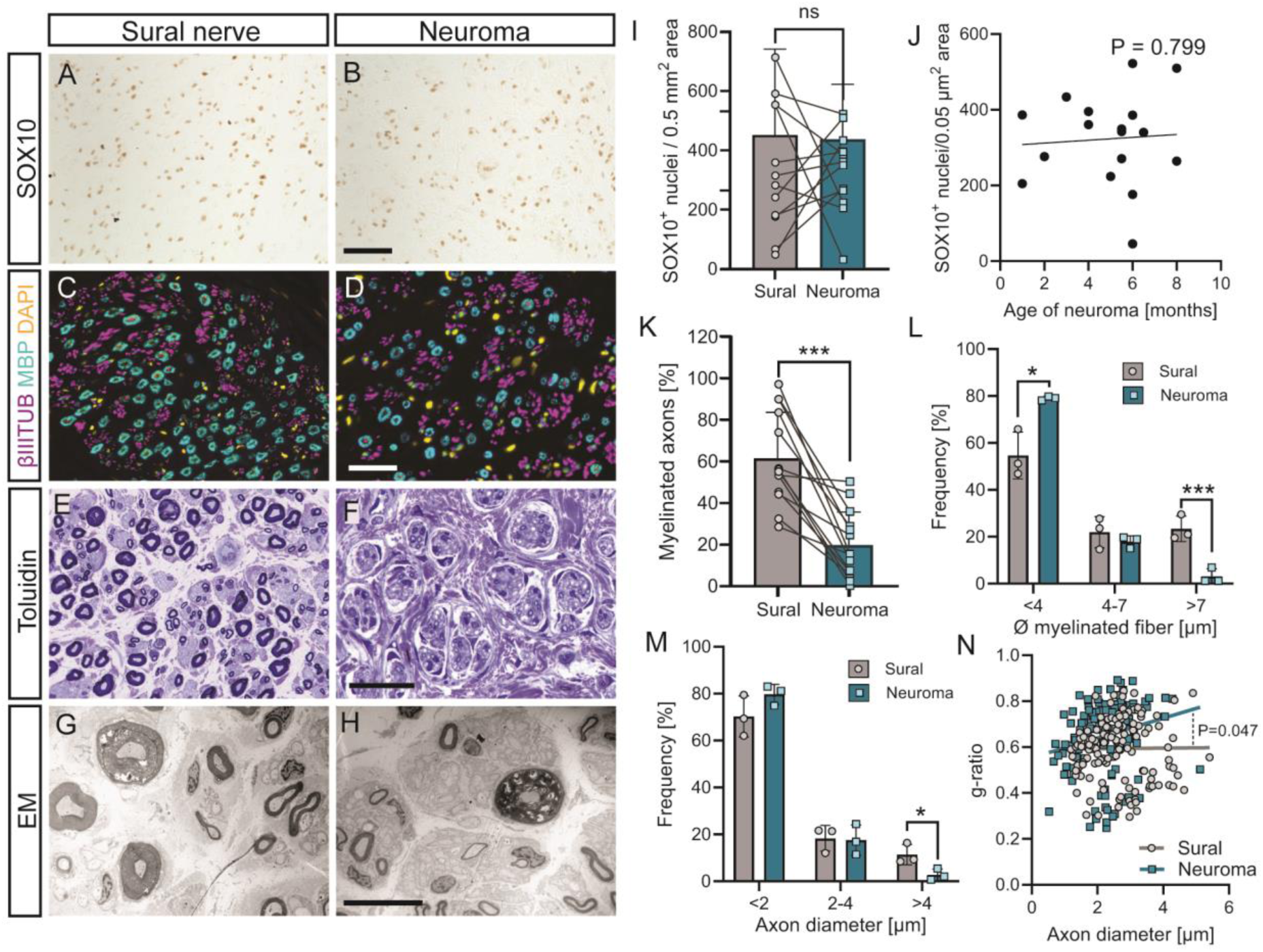
Neuroma formation correlates with reduced numbers of myelinating SCs. (A, B) SOX10 labeled similar SC numbers in sural nerve (A) and neuroma (B). (C, D) Myelinating producing SCs (MBP^+^) were reduced in neuroma (D) compared to sural nerve (C). (E-H) Toluidin (E, F) and EM (G, H) pictures of sural nerve (E, G) and neuroma (F, H) revealed reduced numbers of myelinated axons in neuroma. (I, J) Abundance of SOX10^+^ SCs was similar between sural nerve and neuroma (I) and did not depend on neuroma age (J). (K, L) Quantification depicting reduction of myelinated axons (K) and smaller myelinated fibers (L) in neuroma. (M, N) In neuroma, frequency of large caliber axon was reduced (M) and the g-ratio was increased (N) compared to sural nerve. In (I, K) a line connects sural nerve and neuroma data from the same patient. Each circle in (J, L, M) depicts one patient. Data are presented as mean ± SD. For statistical analysis, Wilcoxon test was performed in (I, K), Pearson’s test in (J), t-test in (L, M), simple linear regression in (N). Statistical significance is provided as *, **, *** reflecting P ≤ 0.05, 0.01 and 0.001, respectively. Scale bar (A-F) = 50 μm, (G, H) = 5 μm

First of all, abundance of SOX10 was investigated (Fig. 3A, B; I). Nuclear abundance of this transcription factor was almost identical between sural nerve (Fig. 3A) and neuroma (Fig. 3B) tissue (see Fig. 3I) and this was not affected by the latency of neuroma formation (Fig. 3J). This suggests no change of total SC numbers (irrespective of maturation status) along the time course of neuroma formation and in relation to an uninjured control nerve. In contrast to total SOX10 positive SCs, numbers of myelin basic protein (MBP) positive SCs were reduced by more than 50% in neuroma (Fig. 3C, E) compared to sural nerve (Fig. 3D, F; quantified in K).

Using toluidine-stained sections and electron microscopy (EM), axonal and myelin ultrastructure was inspected (Fig. 3E-H; L-N). In neuroma (Fig. 3F, H), the percentage of large caliber myelinated axons (Fig. 3L) and the axonal diameter (Fig. 3M) were reduced compared to control nerves (Fig. 3E, G). This was also reflected by higher g-ratios in neuroma compared to sural nerve pointing at thinner myelin sheaths in the former (Fig. 3N).

In summary, myelin producing SCs were diminished in neuroma.

### Neuroma resident SCs express repair and proliferation markers

Data obtained so far show decreased abundance of myelinating SCs in neuroma although total SC number are not altered (Fig. 3). This indicates that SCs might still sustain a non-myelinating repair SC phenotype. To address this, the prototypical repair marker c-JUN was analyzed (Fig. 4). In rodents, c-JUN is functionally required for SCs switching towards a repair mode (Arthur-Farraj *et al*., 2012; Ramesh et al., 2022).

**Figure 4.**
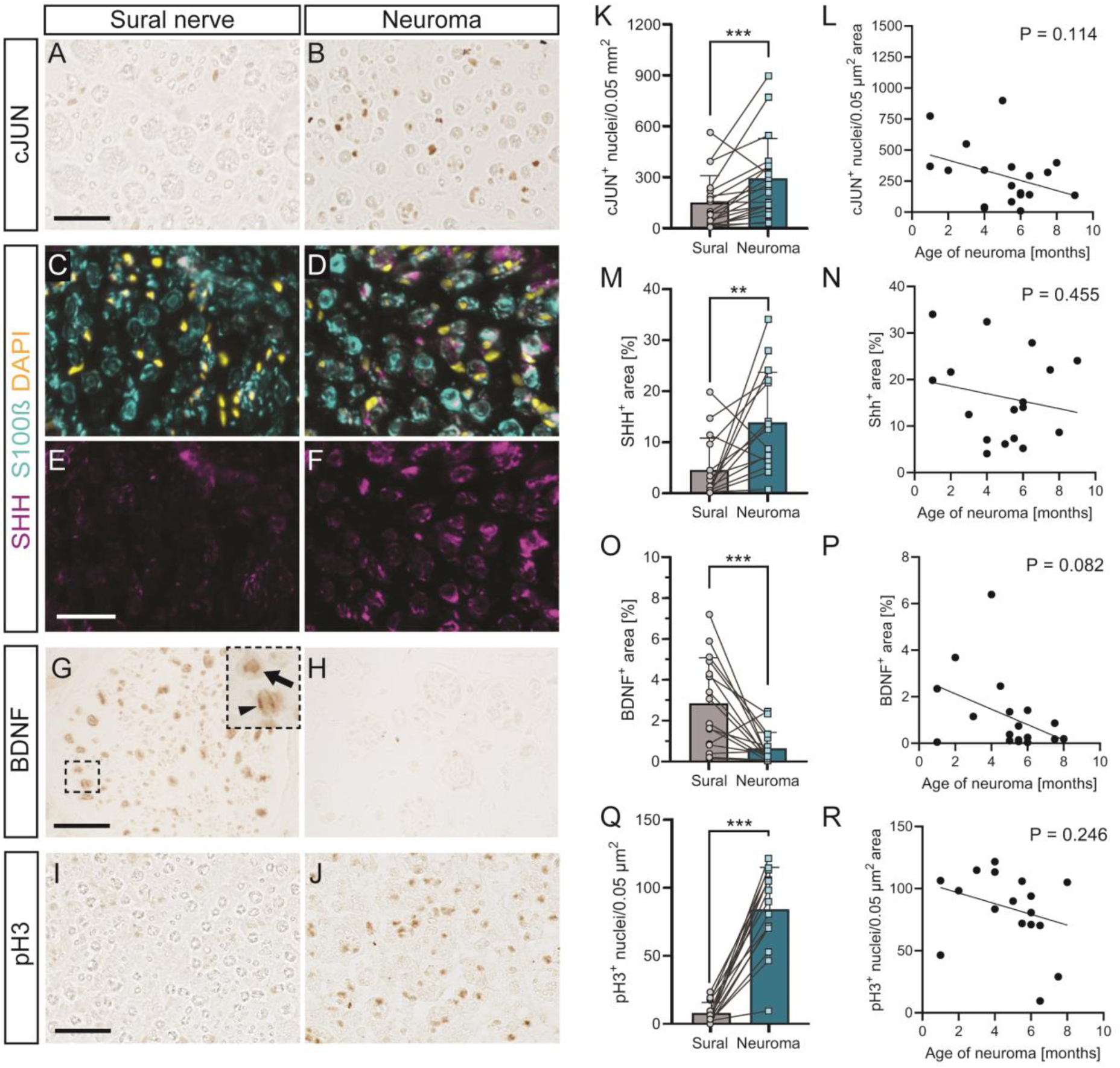
SCs in neuroma sustain a repair phenotype. (A, B) c-JUN was upregulated in neuroma (Fig. 3B) compared to sural nerve (A). (C-F) SHH was expressed in S100 positive SCs. In neuroma (D, F), more SHH^+^ SCs were localized compared to sural nerve (C, E). (G, H) BDNF was more abundant in sural (G) compared to neuroma (H). Insert in (G) shows BDNF localized in axons (arrow) and SC (arrowhead). (I, J) pH3 labels more cells in neuroma (J) in relation to control nerve (I). (K, L) Quantification of c-JUN positive cells between sural nerve and neuroma (K) and correlation with neuroma age (L). (M, N) SHH positive area was higher in neuroma compared to sural nerve (M). No significant correlation with neuroma age was observed although SHH tended to decrease with age (N). (O, P) Decreased PTEN levels in neuroma in relation to sural (O) and a trend towards BDNF downregulation with neuroma age (P). (Q, R) pH3 was ten-fold upregulated in neuroma compared to sural nerve (Q) but numbers remained constant in neuroma over post-injury time (R). In (I, K, M), a line connects sural nerve and neuroma data from the same patient. Each circle in (J, L, N) depicts one patient. Data are presented as mean ± SD. For statistical analysis, Wilcoxon test was performed in (K, M, O, Q), and Pearson’s test in (L, N, P, R). Statistical significance is provided as *, **, *** reflecting P ≤ 0.05, 0.01 and 0.001, respectively. Scale bar (A, B; G, H; I, J) = 50 μm, (C-F) = 25 μm

In neuroma tissue, c-JUN positive SCs (Fig. 4B) were almost doubled compared to uninjured sural nerves (Fig. 4A; K). This suggests a repair phenotype of SCs in neuroma whereas in sural nerve, SCs expectedly remain in a c-JUN negative myelinating state. Notably, with increasing time after injury, c-JUN positive SCs declined in neuroma (Fig. 4L) in line with a previous record (Wilcox *et al*., 2020).

To corroborate this finding, a further marker of repair SCs, sonic hedgehog (SHH; (Arthur-Farraj *et al*., 2012; Jessen and Mirsky, 2021; Ramesh *et al*., 2022), was employed (Fig. 4C-F; M, N). In keeping with c-JUN results, SHH was co-expressed with the SC marker S100β in more SCs in neuroma (Fig. 4D, F) in relation to sural nerve (Fig. 4C, E). Expression of this regeneration promoting morphogen (Moreau and Boucher, 2020) only minimally declines over eight months of neuroma formation (Fig. 4N).

BDNF is a growth factor whose abundance is positively correlated with successful regeneration (McGregor and English, 2018). So far data on BDNF are mostly derived from rodent PNI models and herein we observed decreased BDNF abundance in neuroma (Fig. 4H) compared to intact sural nerves (Fig. 4G; quantified in O, P). BDNF was expressed in both axons (arrow insert Fig. 4G) and SCs (arrowhead; Fig. 4G) in sural nerves.

Finally, strong upregulation of phospho-histone H3 (pH3), a marker labeling proliferative cells, was observed (Fig. 4I, J; Q, R). In fact, pH3 numbers were almost ten-fold upregulated in neuroma (Fig. 4J) compared to control nerve (Fig. 4I; O). Thus, in neuroma tissue several cell types most likely including SCs but also inflammatory cells (see below) appear to proliferate. Numbers of such pH3 positive cells remained rather constant with some tendency to decline along different neuroma ages (Fig. 4R). In summary, our data show that SCs in neuroma express several markers indicative for a repair SC phenotype.

### Neuroma tissue reveals upregulation of pro-regenerative markers

SCs in neuroma express markers indicative of a repair state (Fig. 4). Therefore, we also inspected bona fide pro-regenerative markers to assess the regenerative potential of severed axons and SCs in neuroma (Fig. 5).

**Figure 5.**
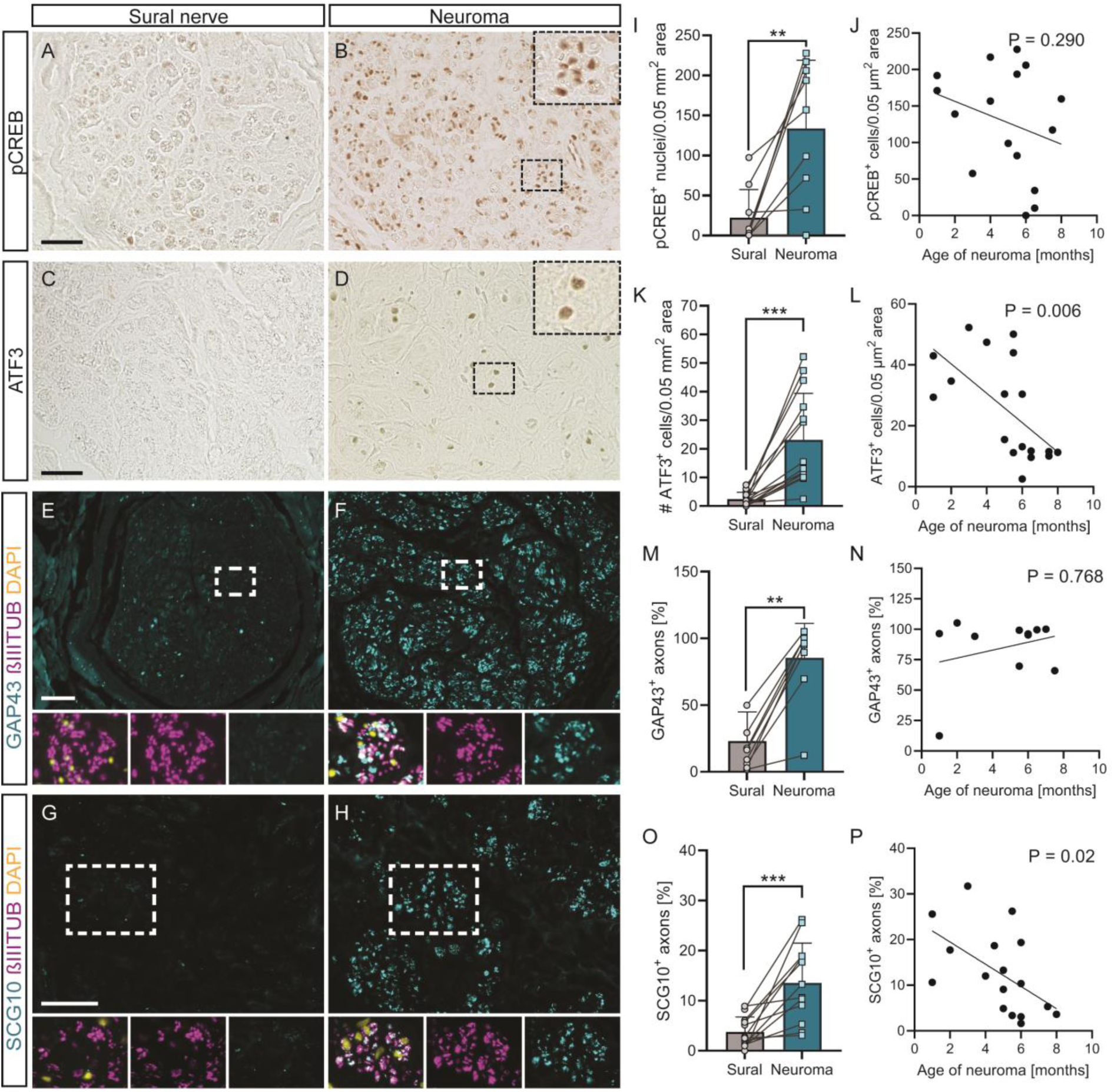
Pro-regenerative markers are induced by axons and SCs in neuroma. (A, B) pCREB was upregulated in neuroma SCs (B) but not sural nerves (A). (C, D) Nuclear ATF3 abundance was increased in neuroma (D) compared to sural nerve (C). (E, F) GAP43 (green) co-localized with axons (magenta). More GAP43 was found in neuroma (F) whereas GAP43 was not expressed in sural control nerves (E). (G, H) SCG10 positive axons (green) co-localized more frequently with axons (magenta) in neuroma (H) compared to sural nerve (G) tissue. (I, J) CREB positive nuclei were more abundant in neuroma (I) with numbers remaining constant in neuroma along different post-injury timepoints (J). (K, L) ATF3 was more abundant in neuroma compared to sural nerve (K) and numbers in neuroma declined over time (L). (M, N) GAP43^+^ axons were enriched in neuroma (M) and remained constant in neuroma tissue over time (N). (O, P) SCG10 was enriched in neuroma axons (O) and decreased with neuroma age (P). In (I, K, M, O), lines connect sural nerves and neuroma data from same patients. Each circle in (J, L, N, P) depicts one patient. Data are presented as mean ± SD. For statistical analysis, Wilcoxon test was performed in (I, K, M, O), and Pearson’s test in (J, L, N, P). Statistical significance is provided as *, **, *** reflecting P ≤ 0.05, 0.01 and 0.001, respectively. Scale bar (A, B, E, F) = 50 μm; (C, D, G, H) = 25 μm

A first marker included was activated CREB (pCREB) enhancing axon regeneration in mouse models and also expressed in rodent SCs (Gao et al., 2004; Stewart, 1995). Herein, we observed pCREB upregulation in neuroma (Fig. 5B). Closer inspection (insert, Fig. 5B) revealed predominant presence of pCREB in SC nuclei (see insert, Fig. 5B). Numbers of such pCREB positive SCs in neuroma persisted independent of neuroma age (Fig. 5F). In contrast, in uninjured sural nerves, no pCREB was present (Fig. 5A; E). Thus, neuroma SCs strongly express an activated pro-regenerative transcription factor.

Similar to pCREB, ATF3 is a transcription factor with regeneration-promoting functions in rodent nerve injury models (Fagoe et al., 2015; Gey et al., 2016; Renthal et al., 2020). Herein, we demonstrate ATF3 upregulation in SCs of neuroma (Fig. 5D) but not sural nerve (Fig. 5C; I). Furthermore, increasing neuroma age negatively correlated with ATF3 abundance (Fig. 5J).

Next, axonally localized markers indicative of enhanced axonal growth were analyzed. This included growth associated protein 43 (GAP43), a protein positively associated with axon regeneration after PNI in rodents (Fu and Gordon, 1997). Similar to pCREB (see above), GAP43 was induced in neuroma tissue (Fig. 5D) compared to sural nerve (Fig. 5C; G) in line with previous reports (Domer *et al*., 2018; Gilmer-Hill et al., 2002). As expected, GAP43 abundance was mainly restricted to axons as demonstrated by co-localization with βIII tubulin (Fig. 5D). Furthermore, GAP43 expression persisted over several months of neuroma development (Fig. 5H).

SCG10, a further marker labeling regenerating axons in rodent PNI (Shin et al., 2014) was analyzed in human injured nerves (Fig. 5G-P). As seen for GAP43, SCG10 abundance was elevated in neuroma resident axons (Fig. 5H) whereas sural nerve axons were negative (Fig. 5G). SCG10 positive axons, however, declined with increasing neuroma age (Fig. 5P).

In summary, SCs and axons in neuroma tissue express several pro-regenerative molecules.

### Neuroma tissue reveals strong presence of inflammatory cells

Inflammatory cells such as macrophages strongly influence the outcome of axon regeneration after nerve injury.

Herein, we tested presence of several macrophage markers including IBA, CD68, CD11c and CD163 in neuroma formation (Fig. 6).

**Figure 6.**
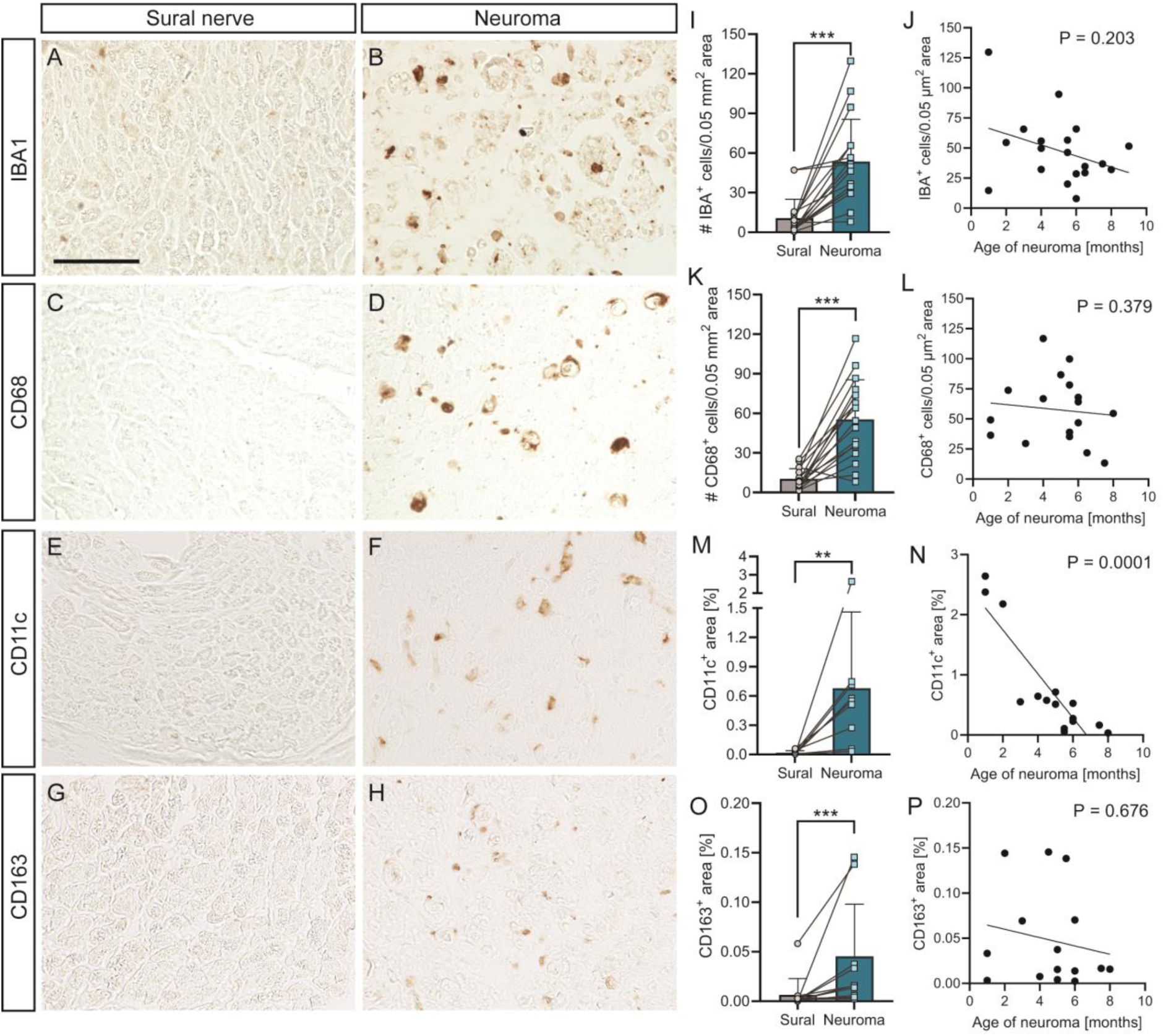
Enhanced presence of macrophages in neuroma. (A, H) In sural nerve, none of the macrophage markers was expressed (A, C, E, G). In contrast, IBA1 (B), CD68 (D), CG11c (F) and CD163 positive macrophages were more strongly present in neuroma. (I, K, M, O) Quantification of macrophage abundance for the different markers. Lines connect sural nerves and neuroma data from same patients. (J, L, N, P) Macrophage abundance is depicted along with the neuroma age. Only CD11c showed a significant reduction in macrophage numbers with increasing neuroma age (N). Each circle in (J, L, N, P) depicts one patient. Data are presented as mean ± SD. For statistical analysis, Wilcoxon test was performed in (I, K, M, O), and Pearson’s test in (J, L, N, P). Statistical significance is provided as *, **, *** reflecting P ≤ 0.05, 0.01 and 0.001, respectively. Scale bar (A-H) = 50 μm

Expectedly, in all uninjured sural nerve samples, none of the inflammation markers was present (Fig. 6A, C, E, G). In contrast, all five inflammation markers were strongly present in neuroma samples (Fig. 6B, D, F, H, quantified in Fig. 6I, K, M, O). Thus, macrophages were upregulated in neuroma, a result which was reported for CD68 before (Donatien et al., 2018; Durrenberger *et al*., 2006; Vora *et al*., 2007). Notably, CD11c (Fig. 6E, F) had the tendency to negatively correlate with neuroma age indicating decreased pro-inflammatory macrophages in older neuroma tissue (Fig. 6N). In contrast, all other markers remained rather constant in neuromas after several post-injury timepoints (Fig. 6J, L, P).

### Cultured neuroma and sural nerve SCs differ in their gene expression profile

*In vivo*, neuroma and sural nerve resident SCs differed in their differentiation state (Figs. 3-6). Next, we addressed whether when taken in to culture and detached from their *in vivo* fibrotic and inflammatory environment, SCs of neuroma and sural nerve might also differ or now show comparable responses to sural nerve SCs (Fig. 7). For this we quantified parameters including SC size, numbers, proliferation, c-JUN expression and gene expression profiles. With regard to cell shape, cultured SOX10-positive SCs of sural nerve (Fig. 7A) and neuroma (Fig. 7B) were morphologically indistinguishable when quantifying area (Fig. 7K). Notably, when taken in culture 100% of all SCs irrespective of whether they were neuroma or sural nerve derived were c-JUN positive (Fig. 7C-F; L). A correlation between c-JUN expression level and non-myelinating SCs has been reported before (Deborde et al., 2022). This is in contrast to lower c-JUN numbers in intact nerves (Fig. 4) but might reflect the injury inflicted on SCs when surgically removed in the patients and by shearing nerves for SC culturing. Furthermore, SC proliferation was also comparable between sural nerve and neuroma SCs (Fig. 7G-J; M) in contrast to *in vivo* (Fig. 4).

**Figure 7.**
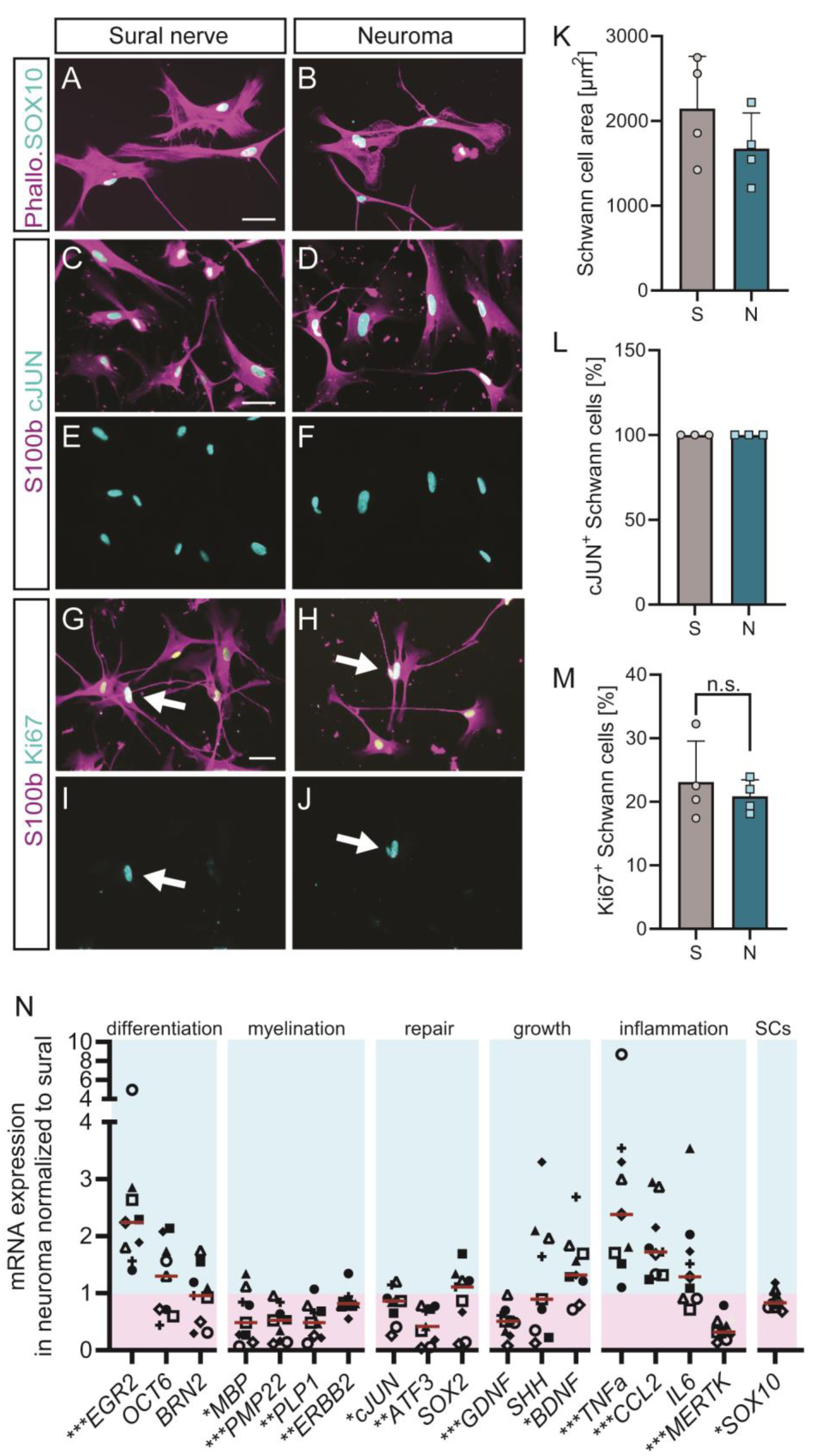
Altered gene expression profiles between cultured neuroma and sural nerve SCs. (A, B) Cultured SOX10 positive SCs of sural nerve (A) and neuroma (B) were morphologically similar. (C-F) All SCs either derived from sural nerve (C, E) or neuroma (D, F) were c-JUN positive. (G-J) The number of Ki67 positive SCs was similar between sural nerve (G, I) and neuroma (H, J). (K) SC area was comparable between sural nerve and neuroma SCs. (L) 100% of neuroma or sural nerve SCs were c-JUN positive. (M) SC proliferation rate was almost identical between sural and neuroma SCs. (N) qPCR data for genes indicated. Levels of sural nerve SC cultures were set to 1 and the values for the corresponding neuroma were calculated accordingly. In (K-N), each geometric symbol depicts one culture derived from one patient. Data in (K-M) are presented as mean ± SD. Red lines in (N) indicate the median. For statistical analysis, Mann-Whitney test was performed in (K-N). Scale bar (A-J) = 50 μm

**Figure 8.**
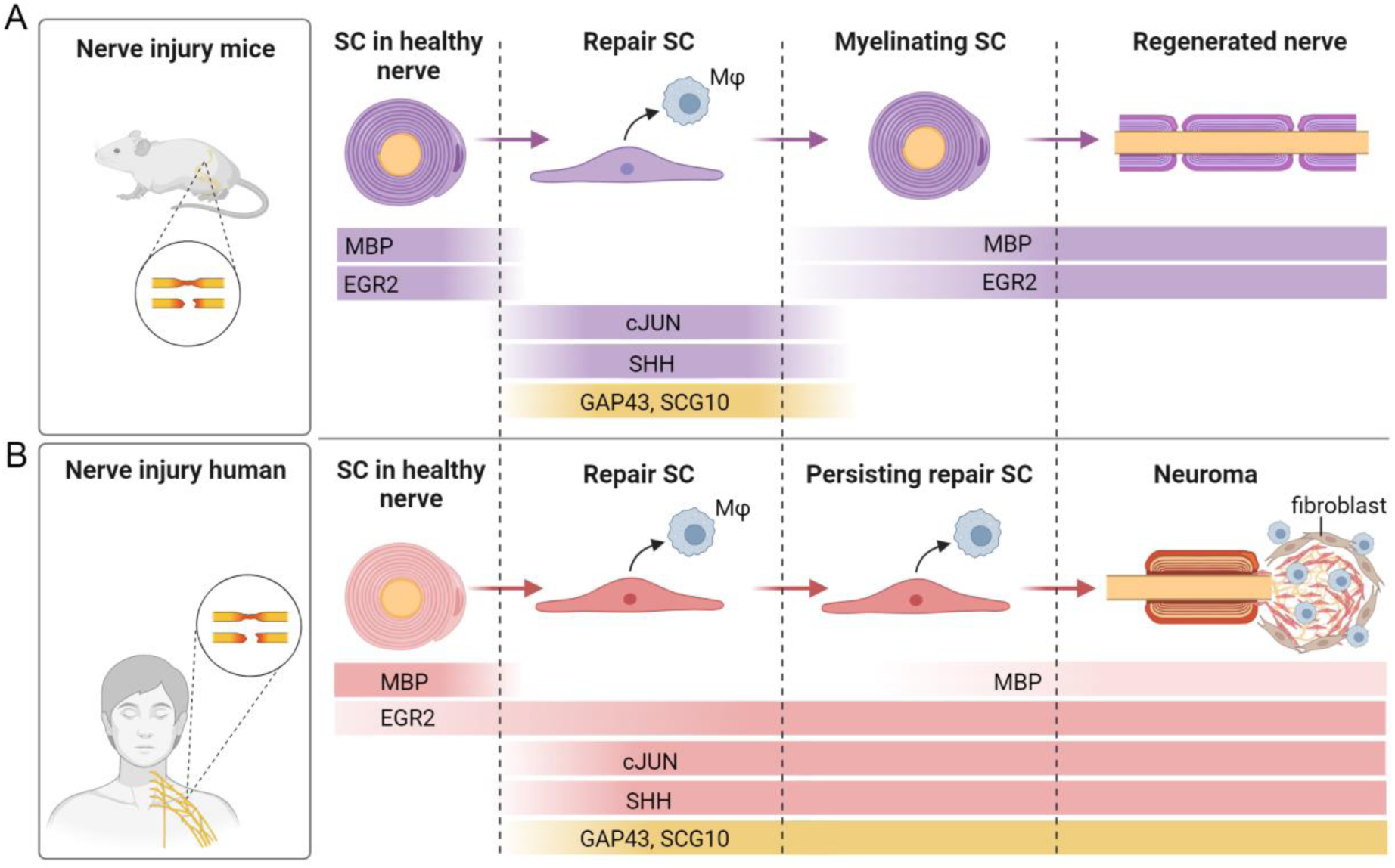
Comparison of rodent and human PNS regeneration. (A) In rodents, PNI induces a switch from myelinating SCs to repair SCs. Repair SCs activate macrophages (Mφ) and express repair markers such as cJUN and SHH. Regenerating axons upregulate markers including GAP43 and SCG10. Eventually, nerve regeneration occurs and SCs start to myelinate again. (B) Summary of findings in human neuroma of this study. Neuroma SCs acquire repair status and express repair markers (cJUN, SHH) similar to rodents. Likewise, axons in neuroma upregulate pro-regenerative molecules (GAP43, SCG10). In contrast to rodents, SCs in human PNI appear to be stalled in repair mode and fail to re-differentiate to myelinating SCs to the same extent. Furthermore, macrophage associated inflammation is persisting. These processes together with fibrosis contribute to neuroma formation at the severed nerve stump and impaired regenerative success.

Thus, morphologically and with regard to proliferation, no differences in cultured neuroma and sural nerve SCs were discernible.

Next, we analyzed expression of markers for differentiation, repair, myelination and e.g. inflammation by qPCR (Fig. 7N). Cultured neuroma SCs revealed higher mRNA levels for differentiation markers such as *EGR2* compared to sural nerve SCs (set to 1; Fig. 7N). Expression of myelin encoding genes including *MBP, PMP22* and *PLP1* was lower in neuroma SC cultures compared to sural nerve SCs (Fig. 6N) in line with *in vivo* findings (Fig. 3). Repair markers such as *cJUN*, despite their similar high abundance on protein level in cultured SCs (Fig. 7C-F), were slightly lower in neuroma SCs (Fig. 7N). Similar results were obtained for two growth factor encoding genes, *GDNF* and *SHH* (Fig. 7N). In contrast, *BDNF* was upregulated in neuroma SCs (Fig. 7N). In keeping with results obtained *in vivo* (Fig. 6), inflammatory genes were more highly abundant in neuroma SC cultures (Fig. 7N).

Taken together, cultured neuroma SCs had higher levels inflammation markers but reduced levels of myelin encoding genes.

## Discussion

### Neuroma SCs adopt repair status but are stalled in this state

In rodent PNI, SCs undergo a defined transition from differentiated myelinating SCs to repair SCs and eventually re-differentiate to myelinating SCs (Bosch-Queralt *et al*., 2023; Jessen and Mirsky, 2019). Transition between these stages is accompanied by SCs expressing myelinating (e.g. myelin genes, *EGR2*) or repair SC (e.g. c-JUN, SHH, BDNF; (Jessen and Mirsky, 2021; Zhang *et al*., 2023)) markers.

In this study we addressed the state of human SCs and asked whether neuroma SCs follow a similar transition (differentiated vs. repair) as described in rodents. It should be kept in mind that neuroma formation is rather unique to human PNI and rodent crush or transection injury models do not perfectly recapitulate characteristic neuroma formation of patients (Toia et al., 2015; Yeoh et al., 2020). Nevertheless, our data together with others (Domer *et al*., 2018; Mahan *et al*., 2019; Wilcox *et al*., 2020) suggest that acquisition of a repair SC state is also induced in neuroma SCs. Hence, similar to repair SCs in rodents, human neuroma SCs have reduced myelin content (Fig. 3) and express the prototypical SC repair marker c-JUN (Fig. 5) in line with previous reports (Wilcox *et al*., 2020). SHH, a repair SC marker known to be upregulated in SCs after rodent PNI (Martinez et al., 2015) was herein described to be also upregulated in neuroma (Fig. 5) as were markers for cell proliferation (Fig. 4). Furthermore, we provide data for two new pro-regenerative markers localized in neuroma so far not described in injured human nerves. ATF3, a regeneration-associated gene (RAG) has established pro-regenerative functions in both injured rodent neurons (Fagoe *et al*., 2015; Gey *et al*., 2016; Renthal *et al*., 2020) and SCs (Hunt et al., 2004; Saito and Dahlin, 2008). In addition, we observed strong CREB activation in neuroma SCs (Fig. 5), a pro-regenerative transcription factor mainly analyzed in injured rodent neurons (Gao *et al*., 2004; Stewart, 1995). In contrast,SOX10 was not changed in neuroma compared to sural nerves in contrast to a previous study (Wilcox *et al*., 2020). This discrepancy might reflect the potential presence of SOX10 in all SC stages covering development, maturation and myelinating SCs as described in rodents (Finzsch et al., 2010). Therefore, the total SC number might remain constant although they shift in number between different maturation stages.

Taken together neuroma SCs acquire several repair markers pointing at a highly conserved post-injury mechanism in the PNS between rodents and humans (see graphical abstract). Nevertheless, this repair status is highest at early post-injury timepoints and declines over time as revealed by our temporal analysis (up to 8 months after injury) for some but not all repair markers (c-JUN, Fig. 4L; ATF3, Fig. 5L). This is congruent with a previous report that likewise demonstrated loss of repair SCs in long-term denervated neuroma tissue (Wilcox *et al*., 2020). Besides SCs acquiring repair phenotypes we also found that injured axons express pro-regenerative markers including GAP43 (in line with (Domer *et al*., 2018)). Furthermore SCG10, a marker frequently used to label regenerating axons in rodent PNS injury models (Shin *et al*., 2014) was also found to be upregulated in human axons in this study (Fig. 5). Interestingly, GAP43 and SCG10 were mainly restricted to small caliber axons (Fig. 5) as also noticed by others (Domer *et al*., 2018). This suggests that such unmyelinated small caliber axons might have strongest regenerative potential. Notably, in rodents SCG10 preferentially labels regenerating sensory rather than motor axons (Shin *et al*., 2014). This suggests for neuroma tissue analyzed herein a stronger upregulation of pro-regenerative markers in the sensory compared to the motor axon population. As noted for repair SC markers above, also some axonal pro-regenerative markers decline over time (SCG10, Fig. 5P). Thus, early post-injury timepoints have highest repair SC presence and also axons in a pro-regenerative state which should result in better regeneration outcomes compared to long-term degenerating nerves. However, given the slow growth rates of severed axons at 1 mm/day (Sulaiman and Gordon, 2013), surgical intervention is often postponed to allow for longer timespans of intrinsic axonal re-growth.

As discussed above, neuroma tissue is positive for both SCs and axons expressing repair and pro-regenerative markers. Why then is neuroma formation typically an obstacle towards successful regeneration? Our data together with other reports (Domer *et al*., 2018; Mahan *et al*., 2019; Tannemaat *et al*., 2007; Wilcox *et al*., 2020) point at several cell-intrinsic but also extrinsic factors in the environment of injured nerves responsible for such a stalling of neuroma SCs in a repair status. Previously, repulsive axon guidance molecules such as semaphorins were found upregulated in neuroma and might contribute towards an axon growth-inhibitory environment (Tannemaat *et al*., 2007). Furthermore, neuroma differs from sural control nerves in neuropeptide abundance (Zochodne et al., 1997). Growth factor supply is an important factor for enhancing axonal growth (McGregor and English, 2018). Herein, BDNF was found downregulated in neuroma tissue compared to control nerve and therefore might contribute to failed axonal re-growth. In addition, inflammation and fibrotic scar tissue are considered extrinsic factors impeding on axon regeneration (Durrenberger *et al*., 2006; Karsy *et al*., 2018; Mahan *et al*., 2019). In agreement, we found strong infiltration of neuroma tissue by macrophages in contrast to un-injured nerves which were devoid of such inflammatory cells (Fig. 6; see graphical abstract). Of note, CD11c macrophages are pro-angiogenic and induce neovascularization (Droho et al., 2023) which would be in accordance with the higher vascularization observed in neuroma (Fig. 2). Furthermore, CD163 positive macrophages are pro-fibrotic (Kawanaka et al., 2023). This later finding is in agreement with more fibrotic cells observed in neuroma (Fig. 2).

These extrinsic growth obstacles suggest that the environment encountered by SCs plays a major role in the regeneration outcome. Thus, when removed from their inhibitory environment, neuroma SCs might more readily fulfil pro-regenerative functions. To test this hypothesis, SCs from neuroma and control nerve were taken in culture to monitor responses of individual SCs (Fig. 7). Interestingly, when quantifying numbers and morphological parameters, neuroma and control SCs were undistinguishable (Fig. 7). This indicates that when neuroma SCs are liberated from a growth-inhibitory environment they have similar adhesive, growth and proliferation potential as uninjured human SCs. Nevertheless, it was interesting to see that neuroma SCs retain a gene expression signature which in some aspects is similar to their *in vivo* situation (Fig. 7). This included reduced myelin and elevated inflammatory gene expression when compared to cultured sural control SCs. In the future, it will be interesting to identify molecules which ensure the *in vivo* neuroma SC identity also in culture. Particularly epigenetic mechanisms might be involved in retaining *in vivo* SC properties also in culture.

### Technical considerations and limitations of this study

Work with human nerve samples has limitations compared to rodent models. First of all, human nerve samples obtained only allow to analyze responses of nerve-resident cells or compartments including SCs, axons and other cells (e.g. endothelial cells, fibroblasts, macrophages). However, this study precludes to investigate responses of neuronal cell bodies (e.g. transcription) localized in e.g. dorsal root ganglia (DRG) which is typically done in rodent PNI. Secondly, nerve and neuroma tissue obtained was highly heterogenous with regard to patient age, post-injury latency time and patient medication which is a source of data variability (see Table 1). A further limitation was the nerve types used for comparison. We compared responses of neuroma mostly derived from the brachial plexus which consists of both sensory and motor fibers with purely sensory sural nerves which lacks motor fibers. Here, it also has to be kept in mind that control sural nerves were not completely uninjured. Since they were surgically removed, injury (i.e. cut with a scalpel) was also introduced and some injury responses might occur *ex vivo* until samples were frozen (taking 10-30 minutes). Finally, n numbers for neuroma isolated shortly after injury (1-5 months) were rather low (n=4-6) compared to longer latency periods with higher n numbers (see Table 1). However, this distribution was in line with other studies (Wilcox *et al*., 2020) and reflects the fact that PNI surgery is typically performed at later post injury time points (> 6 months) to allow for maximal spontaneous axon re-growth.

## Conclusions

Herein we show that SCs in injured human neuroma nerve tissue adopt several repair markers and axons express pro-regenerative molecules which however decline with progressing neuroma age. This suggests a cross-species conservation of SCs switching after injury from a myelinating to a repair phenotype. However, unlike in rodents, SCs in human neuroma seem to be stalled in a repair phenotype and fail to support axon regeneration and eventually re-differentiate in myelinating SCs again. The latter might be inflicted by cell intrinsic (e.g. growth factor deprivation) or extrinsic factors including inflammation and tissue fibrosis. Interestingly, when neuroma SCs are grown in culture they have almost identical growth properties compared to uninjured SCs. Still, they retain a molecular profile of enhanced inflammatory and lower myelin gene expression.

## List of abbreviations

ATF3: activating transcription factor 3
BDNF: brain derived neurotrophic factor
CREB: cAMP responsive element binding protein
DAB3: 3’-diaminobenzidine
DRG: dorsal root ganglia
EGR2: early growth response 2
GAP43: growth associated protein 43
GDNF: glial derived neurotrophic factor
HPRT: hypoxanthine phosphoribosyltransferase 1
iDISCO: 3D Imaging of Solvent-cleared Organs
MBP: myelin basic protein
NIC: neuroma in continuity
pH3: phospho histone H3
PNI: peripheral nerve injury
PNS: peripheral nervous system
RAG: regeneration associated gene
SC: Schwann cell
SHH: sonic hedgehog
SOX10: SRY-box transcription factor 10

## Declarations

### Ethics approval and consent to participate

All donors had provided written informed consent for the use of biopsy material and of clinical and genetic information for research purposes.

### Availability of data and material

The datasets supporting the conclusions of this article are included within the article.

### Authors’ contributions

SD, SMzR and BK planned the experiments. SD, JS, AB and CK performed the experiments and evaluated all data. MP and GA contributed patient samples. SD, SMzR and BK prepared figures and drafted the manuscript. All authors read and approved the manuscript.

## Acknowledgements

The authors thank Daniela Sinske for technical support and Nadine Tietz for excellent work during her lab rotation. The graphical abstract was generated with BioRender.

